# SiGMoiD: A super-statistical generative model for binary data

**DOI:** 10.1101/2020.10.14.338277

**Authors:** Xiaochuan Zhao, Germán Plata, Purushottam D. Dixit

## Abstract

In modern computational biology, there is great interest in building probabilistic models to describe collections of a large number of co-varying binary variables. However, current approaches to build generative models rely on modelers’ identification of constraints and are computationally expensive to infer when the number of variables is large (*N* ∼100). Here, we address both these issues with **S**uper-stat**i**stical **G**enerative **Mo**del for b**i**nary **D**ata (SiGMoiD). SiGMoiD is a maximum entropy-based framework where we imagine the data as arising from super-statistical system; individual binary variables in a given sample are coupled to the same ‘bath’ whose intensive variables vary from sample to sample. Importantly, unlike standard maximum entropy approaches where modeler specifies the constraints, the SiGMoiD algorithm infers them directly from the data. Notably, unlike current approaches, SiGMoiD allows to model collections of a very large number (*N* > 1000) of binary variables. Finally, SiGMoiD offers a reduced dimensional description of the data, allowing us to identify clusters of similar data points as well as binary variables. We illustrate the versatility of SiGMoiD using several datasets spanning several time- and length-scales.

## Introduction

In recent years, there has been great interest in modeling the statistics of a large number of co-varying binary variables. Significant examples can be found in genomics, where the presence or absence of thousands of genes across microbial genomes dictate phenotypes^1^ or co-varying mutations in protein sequences dictate their fitness^2^, in microbial ecology, where the presence or absence of species across microbiomes is indicative of direct metabolic interactions^3^, or in neuroscience, where hundreds of thousands neurons spike in a correlated manner in order to carry out organism-level tasks^4^. Our ability to collect these high dimensional data has improved substantially in the last decade. Yet, estimating the frequency of occurrence of every possible binary configuration from available samples is not possible for any reasonably sized collection; a system with *N* co-varying binary variables has 2^*N*^ possible configurations and the number of samples available is typically orders of magnitude lower than the number of configurations.

At the same time, given the complexity of interactions, it is infeasible to build bottom-up mechanistic models to describe these systems. A popular alternative is to derive approximate top-down probabilistic models and train those models on the data. Over the past two decades, the maximum entropy (max ent) method^5^ has emerged as perhaps the only candidate for building approximate generative models across a variety of contexts^6-9^. Here, one computes user-specified lower order statistics from the samples and seeks the maximum entropy Gibbs-Boltzmann distribution consistent with these data-driven constraints. A significant conceptual advantage of max ent is that it seeks unbiased models except for the constraints imposed by the modeler^5,l0^. On the flip side, max ent requires the modeler to *a priori* identify problem-appropriate constraints, which may or may not be known. Current max ent approaches have practical limitations as well; inference of model parameters requires costly Markov chain Monte Carlo (MCMC) simulations that prohibits models of *N* > 100 co-varying binary variables^11^.

In order to study covariation in a large number of binary variables in a constraint-agnostic and numerically efficient manner, we propose a novel dimensionality reduction framework inspired from physics; **S**uper-statistical **G**enerative **Mo**del for b**i**nary **D**ata (SiGMoiD). SiGMoiD is a generalization of the max ent approach and has several salient features that distinguish it from the state-of-the-art models of binary variables. <u>First</u>, SiGMoiD does not require the modeler to choose the max ent constraints; the inference procedure learns them directly from the data. <u>Second</u>, SiGMoiD is a dimensionality reduction method. The lower dimensional embedding of the space of data points and the space of binary variables can be used to cluster “similar” data points as well as binary variables. <u>Third</u>, SiGMoiD is a generative model, it allows us to assign probabilities to unseen data and to generate *in silico* samples that capture essential co-variation features of the data. <u>Finally</u>, the inference in SiGMoiD is significantly faster than the current approaches, allowing us to model collective variation in *N* > 1000 binary variables; a system size that is an order of magnitude higher than the current approaches. Below, we first sketch the outline of SiGMoiD and then illustrate it utility by applying it to several data sets.

## Results

### The model

We assume that individual samples comprise *N* binary variables {*σ*_*i*_} (*i ∈* [1, *N*]) that take values 0 or 1. Let us denote by *π*_*i*_ the probability that *σ*_*i*_ = 1 and by ***π*** the vector of probabilities ***π*** = {*π*_1_, *π*_2_,…, *π*_*N*_}. To motivate our model framework (Fig. 1), we imagine the following physical process: each binary variable in the collection of *N* variables is interacting with the same bath that can exchange *K* types of extensive variables (energies). The *k*^*th*^ type of energy for each binary variable in the state when it is active (*σ*_*i*_ = 1) is *E*_*ki*_ and zero when it is inactive (*σ*_*i*_ = 0) (denoted collectively by *E*. Under these circumstances, the probability of the *i*^*th*^ binary variable is equal to 1 is given by the Gibbs-oltzmann distribution^12^:

**Figure 1.**
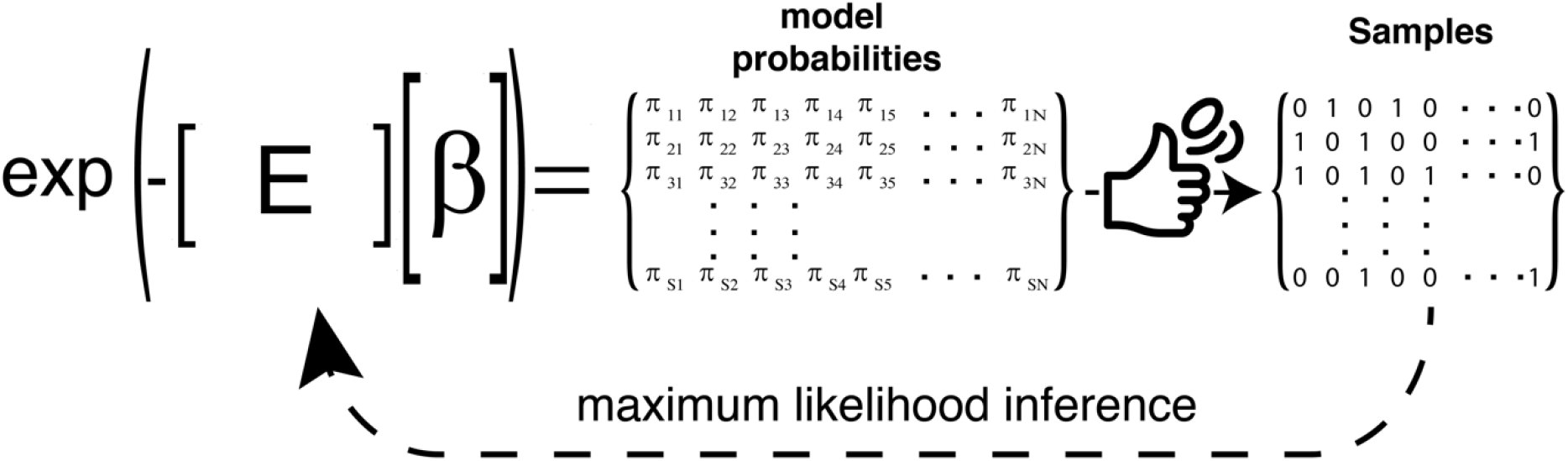
Schematic of the SigMoiD approach. Probabilities *π*_*is*_ for variables *i* in samples s are generated according to a Gibbs-Boltzmann distribution with energies ***E*** and latents *β*. The observed data (samples) is a collection of Bernoulli trials based on the model probabilities. SiGMoiD infers the parameters ***E*** and *β* using maximum likelihood inference.

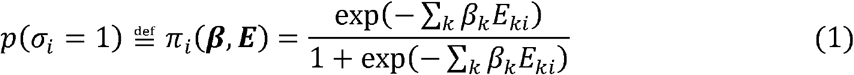

In Eq. 1,***β***={*β*_1_, *β*_2_,…, *β*_*K*_} are the intensive variables. The probabilities in Eq. 1 are the maximum entropy probability distributions when averages ⟨ *E*_*ki*_⟩ (*k ∈* [1, *K*]) of the *K* types of energies are specified for each variable (*i ∈* [1, *N*]).

We set up the model such that the intensive variables vary from sample to sample but are shared across all binary variables in a given sample. In contrast, the energies vary from variable to variable but are shared across all samples. Let us consider that we are given *K* samples {*σ*_*is*_}(*i ∈* [1, *N*]) (*s ∈* [1, *S*])of the binary variables. From these samples, we infer sample-specific intensive variables ***β***_*s*_ and sample-independent energies ***E*** from the data. To that end, we take a maximum likelihood approach. We write the log-likelihood of the data given the parameters:

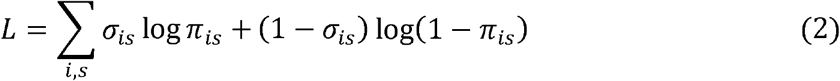

The log-likelihood can be maximized to determine the parameters using gradient ascent. The gradients are given by:

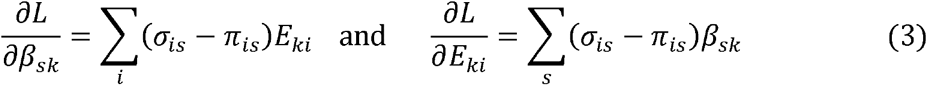

SiGMoiD has several salient features. First, similar to other non-linear dimensionality reduction methods, if *K« N*, SiGMoiD offers a reduced description of the data; the *K* dimensional vectors ***β***_*s*_ embed the *N* dimension data point ***σ***_*s*_ in a *K« N* dimensional space. In addition, since SiGMoiD is a fully probabilistic approach, it can also be used as a generative model. Random samples can be generated as follows. We first select a random set of intensive variables ***β***_*s*_, evaluate the probabilities ***π***_*s*_ and sample random variables ***σ*** as Bernoulli variables using those probabilities. Finally, SiGMoiD also allows us to evaluate the probability of a new set of binary variables ***σ*** given the other observations. Specifically, the probability is

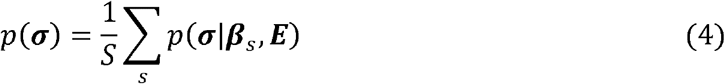

where

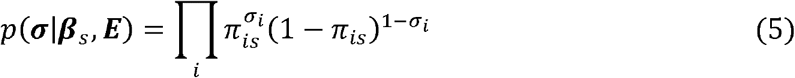

is the probability of observing the binary variables ***σ*** when the intensive variables are fixed at ***β***_*s*_. Below, using several examples, we show the utility of SiGMoiD across a range of applications.

### Accuracy of SiGMoiD as a probabilistic model: modeling the collective firing of neurons

Before illustrating SiGMoiD using larger data sets, we first show a comparison between SiGMoiD and the standard approach to model binary variables; a max ent model. We use a previously collected data set measuring the collective firing of 160 retinal neurons for the duration of a movie that lasted 19 seconds^13,14^ (see Supplementary Information). We note that inference of a max ent model for the collective firing of all 160 neurons is currently computationally prohibitive. We chose the 15 most active neurons in the data (15 highest firing propensities) to illustrate our approach. First, we inferred a max ent model from the data that constrained mean firing rates and pairwise correlations. The max ent model describes the probability of any configuration ***σ*** as:

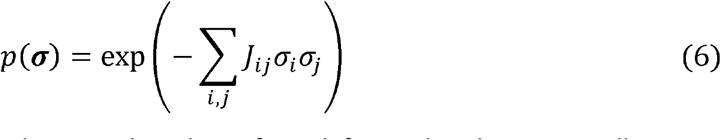

In Eq. 6, *J*_*ij*_ are coupling constants that need to be inferred from the data, typically using gradient descent^9^. Given that there are only 2^15^∼3×10^4^ states for 15 neurons, we could estimate model predictions and therefore the coupling constants by an exhaustive force summation over all possible states without resorting to MCMC simulations. This minimized the errors in max ent inference that arise due to inaccuracies in MCMC-based estimates of average firing rates and neuron-neuron correlations.

In Fig. 2, we compare the two models. The max ent model has 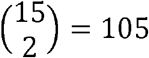 neuron-specific parameters. To match that number, we choose *K*= 7 in SiGMoiD. In panel (a), we show a comparison between the raw probabilities of individual configurations obtained from data (x-axis) to model predicted probabilities (y-axis, red: max ent, blue: SiGMoiD). It is clear that the SiGMoiD model has smaller error compared to the max ent model (mean absolute error 7.4 × 10^−6^ vs 1.4 × 10^−5^). In panel (b), we plot the probability *p*(*n*) that *n* neurons fire at the same time as observed in the data (black), predicted using SiGMoiD (blue), and using the max ent model (red). Here too, the SiGMoiD model performs well when capturing the likelihood. In panels (c) and (d), we plot the three-body correlations ⟨*δ*σ_*i*_ *δ*σ_*j*_*δ*σ_*k*_⟩ as observed in the data (x-axis) and as predicted by the model (y-axis, SiGMoiD, panel (c), max ent, panel (d)). Both models capture the three body correlations with reasonable accuracy; the mean absolute error is 9.7 × 10^−4^ vs 1.4 × 10^−3^ for the SiGMoiD and the max ent model respectively. This analysis illustrates that the SiGMoiD is better than the max ent based model at capturing the data and making predictions.

**Figure 2.**
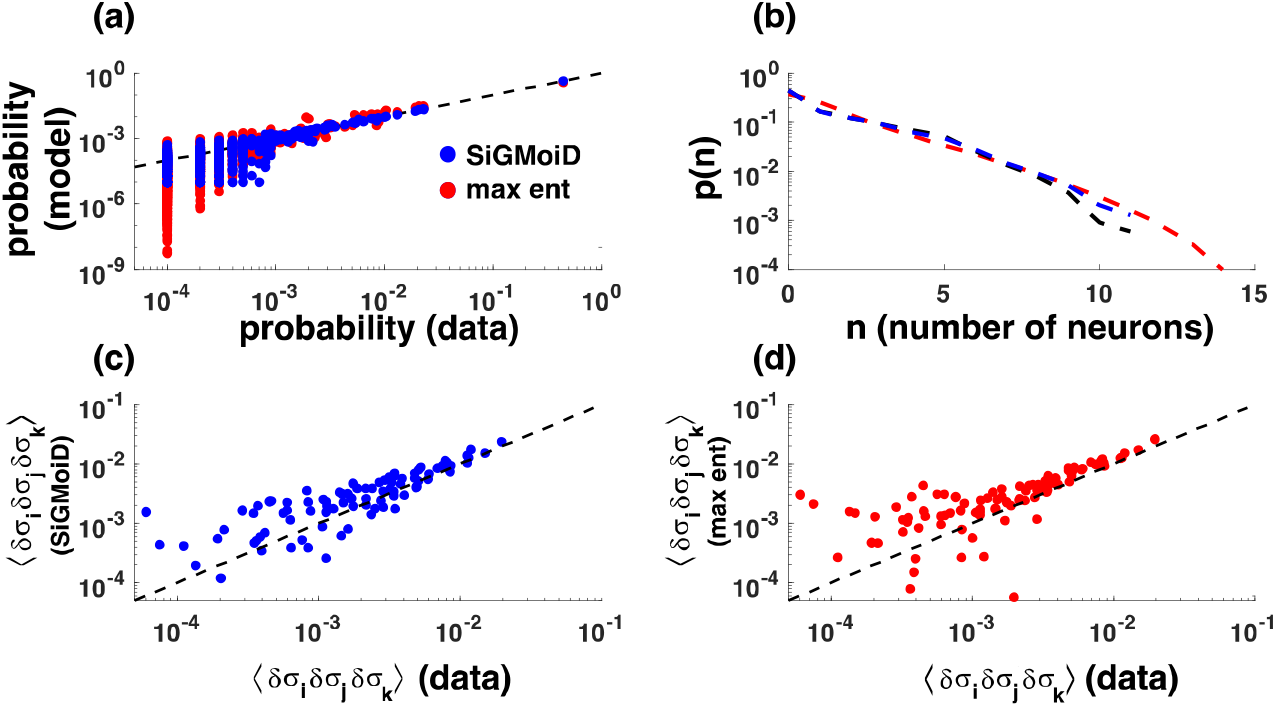
Comparison of SiGMoiD with max ent modeling. **(a)** the probabilities of individual configurations estimated from the data (x-axis) and from the two models (y-axis), (red: max ent, blue: SiGMoiD) **(b)** the probability *p*(*n*) that *n* neurons fire in any given configuration as estimated from data (black), the max ent model (red), and SiGMoiD (blue), **(c)** and **(d)** comparison between three variable correlations ⟨*δσ*_*i*_ *δσ*_*j*_*δσ*_*k*_⟩ estimated from data (x-axis) and those using the models (y-axis).

### Inference of interactions from bacterial co-occurrences using SiGMoiD

Gut microbiomes are complex ecosystems whose statistical properties have received significant attention in the last couple of years^15,16^. Gut bacteria live in species-rich communities where they compete for nutrients and also exchange metabolites with each other. Describing these interactions is critical to map the ecological networks of gut microbiomes and identify targets for controlling microbial communities^17^. However, many of the direct metabolic interactions between gut microbes are likely to occur at a micron length scale^3^. Therefore, it is infeasible to infer these interactions from macroscopic, community-wide abundance co-variation.

To address this issue, Sheth et al.^3^ recently probed the spatial organization of the gut microbiome at the micron length scale, allowing them to capture putative direct interactions between bacteria. In these experiments, Sheth et al.^3^ fractionated mice guts into particles with a median diameter of 30 *µm* and quantified the membership of ∼ 350 operational taxonomic units (OTUs) across ∼ 1500 particles. However, given that co-occurrences are transitive (if A interacts and co-occurs with B, and B interacts and co-occurs with C, then A co-occurs with C even in the absence of interactions), it is not possible to use simple co-occurrence calculations to identify putative pairs of directly interacting OTUs^18^.

Given that SiGMoiD can directly model occurrence of individual OTUs across particles, it can be used to identify clusters of OTUs that co-vary across particles as well as clusters of particles that show specific OTU occurrence profiles. We therefore analyzed the data collected by Sheth et al.^3^ using SiGMoiD (see Supplementary Information). Each particle was characterized by a binary vector representing the OTUs present in that particle. It is evident that SiGMoiD will fit the data better as the number of components *K* increases. Unlike the neuron samples which were correlated in time and across different trials, the microbiome particles are likely to be closer to statistical independence. Therefore, we can use information theory-based criteria to select the optimal *K* that fits the data but avoids overfitting. In SI Figure 1, we show the Akaike information criteria (AIC) vs. *K* for the OTU data. The model picks out *K* = 8 as the optimal value which we use in further analysis.

The number of species found in each particle, a gross descriptor of the complexity of the community^7^, varies substantially from particle to particle. As shown in Fig. 3, generative modeling of the particles using SiGMoiD accurately captures this quantifier of ecological complexity. In panel (a) of Fig. 3 we show the probability of co-occurrence of multiple OTUs in any community as observed in the data (black circles) and as predicted by SiGMoiD (blue line). Notably, the observed distribution of OTUs in different particles is markedly different than the one expected based on the occurrence frequency of individual OTUs (red line). This suggests that SiGMoiD can accurately capture the interactions between OTUs, which in turn allows it to predict the co-occurrence distribution.

**Figure 3.**
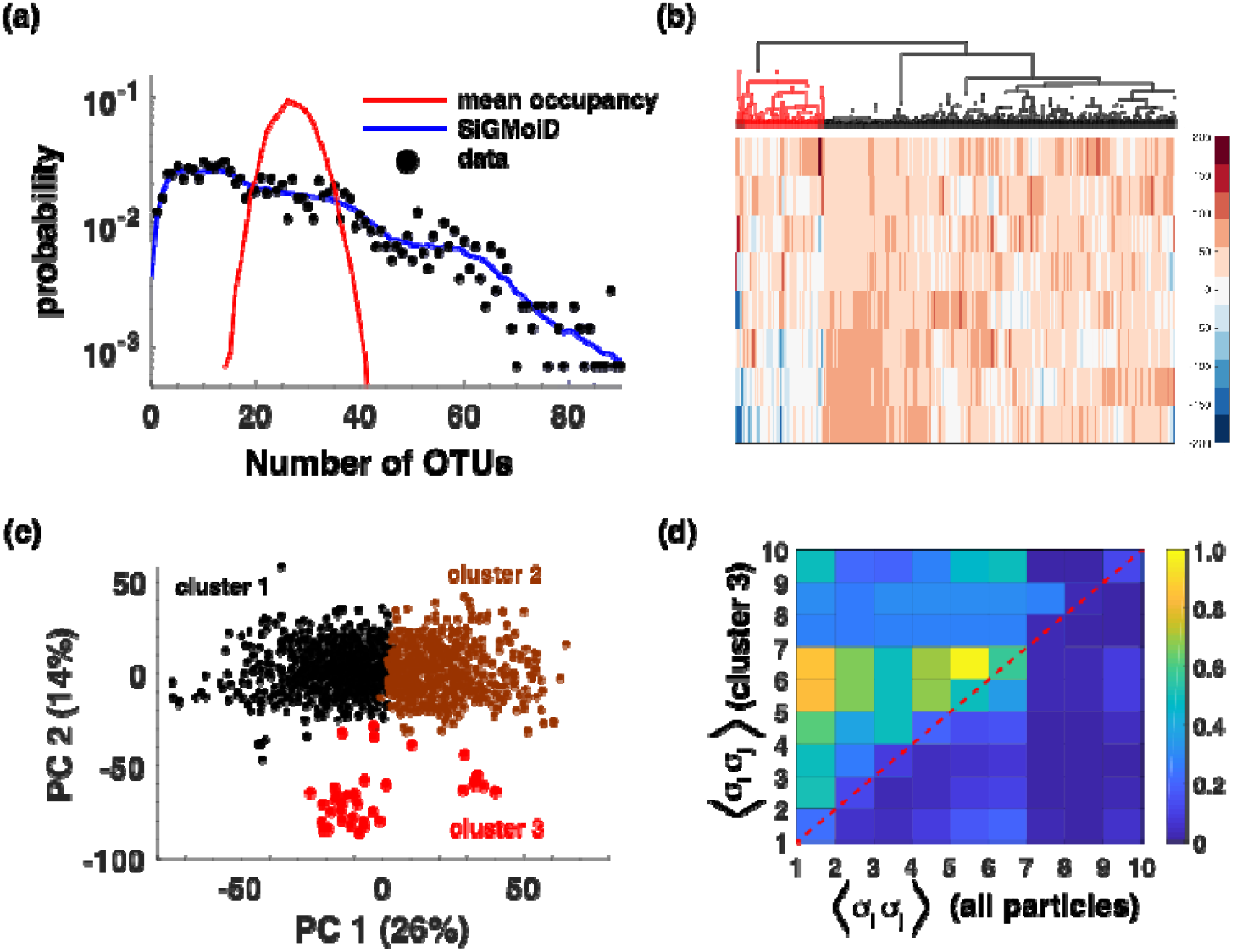
SiGMoiD models bacterial co-occurrences and interactions. (**a**) the probabilities of co-occurrence of multiple OTUs in a single particle. Black circles represent the data, the blue line and shaded blue region represents the SiGMoiD predictions and standard deviations around the predictions, and the red line represents a prediction based on mean occupancies of OTUs. (**b**) Clustergram showing similarity in features between OTUs. The identified outgroup is marked with a dashed line. (**c**) PCA of 3 clusters identified using particle-specific low dimensional embeddings. (d) (upper half) Co-occurrence frequencies of 10 OTUs whose occurrence frequency was most significantly different in cluster 3 compared to the the baseline co-occurrence frequencies of the same OTUs (bottom half).

In fact, SiGMoiD can be used to identify specific bacteria with similar occurrence profiles across particles. SiGMoiD characterizes each binary variable (here, OTU presence/absence) using a *K* dimensional vector of energies. OTUs with similar energy profiles will have similar co-occurrence profiles as well. Therefore, the energy vector can be used to identify clusters of co-occurring OTUs. SiGMoiD-based clustering of OTUs is a more direct way of identifying clusters by relying on inferred inherent properties of the OTUs (the energies) rather than their co-occurrence profiles. Panel (b) of Fig. 3 shows a hierarchical clustering plot of all OTUs using SiGMoiD-inferred energies. Among the several identified clusters, we focus on the cluster of 69 OTUs highlighted in the figure. The gut microbiome of mice is dominated by OTUs belonging to the family *Lachnospiraceae;* ∼ 53% of all the OTUs in the analyzed data belonged to this family. However, these OTUs are not equally distributed across the particles. The cluster highlighted in the figure is statistically significantly enriched with the family *Lachnospiraceae* (46 out of 69, single tailed hypergeometric distribution p-value 0.009). Notably, the OTUs belonging to this cluster had predominantly positive correlations across different particles; 2330 out of the 2346 unique pairs had a positive correlation with 92% of pairs with a p-value less than 10^−2^ (84% of pairs with a p-value less than 10^−4^ and 74% of pairs with a p-value less than 10^−6^). In comparison, only 50% of unique pairs from other OTUs had a positive correlation and only 23% of those correlations had a p-value less than 0.01. These analyses suggest that SiGMoiD-based features can identify clusters of OTUs that significantly co-occur in a given ecology. There are two types of metabolic interactions between bacteria that lead to co-occurrence in an ecosystem^19^, especially at the micron length scale^3^. Genetically related bacteria tend to co-occur because they have similar metabolic networks and can compete for the same resources. In contrast, genetically dissimilar bacteria have different metabolic networks and can cross-feed each other; one species utilizing the metabolic byproducts of another. Therefore, this cluster likely represents the co-occurrence of multiple species in the *Lachnospiraceae* family that compete with each other for the same resources in the mouse gut.

In addition to identifying OTUs that have similar occurrence profiles across communities, SiGMoiD can also be used to identify communities that have similar OTU occurrence profiles. SiGMoiD embeds each high dimensional binary sample in a much lower dimensional space of sample-specific ***β*** vectors. Using K-means clustering of sample-specific ***β*** vectors, we identified 3 clusters of particles (SI Figure 1). Principal component analysis (PCA)-based visualization of the particles clearly shows the three identified clusters (Fig. 3, panel c). Notably, several specific OTUs were co-present with much higher occurrence frequencies in the identified small cluster (cluster 3, comprising 47 particles). In panel (d) of Fig. 3, we compare the pairwise co-occurrence frequency of 10 OTUs whose occurrence frequency was identified to be most significantly different between particles in cluster 3 compared to the baseline using a hypergeometric test (SI Table 1). It is clear that compared to the baseline co-occurrence frequency (sub-diagonal half of Fig. 3 (d)), the pairwise co-occurrence frequencies of the 10 OTUs are significantly elevated in the communities in cluster 3 compared to the rest of the communities. These analyses show that SiGMoiD can also identify specific communities that comprise strongly co-occurring bacteria that differentiate them from other communities. These significant clusters can potentially be investigated for direct co-operative or competitive interactions, as well as their association with distinct regions of the gut. Importantly, clusters with these tightly correlated species were not detected when we applied the same clustering approach to the full microbiome data collected across different particles (SI Figure 2).

### Identifying missing metabolic reactions using SiGMoiD

The metabolic repertoire of microorganisms enables them to convert nutrients into biomass and energy and underlies phenotypic traits central to their ecosystem roles. E.g. microbial fermentation in the gut and its impact on human health^20^ or methane production in animal agriculture or wetland ecosystems^21,22^.

Genome sequencing and annotation methods have enabled the identification of metabolic transformations that individual microbes can potentially carry out through the reconstruction of their metabolic networks^23^. Metagenomics sequencing on the other hand, has allowed the study of the genomes and metabolic properties of microorganisms in microbiomes of interest which have not yet been cultured and characterized. Nevertheless, due to the complexity of these microbial communities, it is often not possible to determine the full genomic content of most members of a given microbiome. Therefore, beyond a few highly abundant microbes whose genomes can be inferred to a reasonable degree of completeness from metagenomics data, the metabolic capabilities of many microbes of interest can only be partially assessed through metagenome annotation and binning methods. Here we show that SiGMoiD can be used to infer missing reactions in the metabolic repertoire of incompletely sequenced genomes, as are often produced by metagenome assembly and binning pipelines.

To that end, we downloaded genome-scale metabolic reconstructions for ∼4000 bacteria from Kbase^24^ generated with the ModelSeed pipeline^25^ (see Supplementary Information). Often, intercompartmental transport reactions and other reactions are added to metabolic reconstructions to ensure mass balance and viability of biomass production without a clear identification of the genes that may carry out these reactions. To avoid biasing our approach towards or against these *ad hoc* additions, we only retained those reactions that were assigned to one or more genes in the reconstructions. This resulted in a total of ∼3300 reactions across all bacterial metabolic reaction sets. We randomly selected 400 bacteria as a test set to quantify the accuracy of our predictions. We inferred SiGMoiD parameters on the rest of the bacteria and used the inferred parameters to identify missing reactions in the test dataset as follows.

First, for each bacteria in the testing set, we removed a fixed fraction of reactions ranging from 10% to 90% to simulate incomplete genome coverage from metagenomic sequencing. Next, we used SiGMoiD and the known reactions for each bacteria in the testing set to predict the missing reactions. To that end, for any given *K*, we identified groups of reactions with similar occurrence profiles across the bacterial world by clustering their corresponding *E*_*ki*_ values obtained from the training data using hierarchical clustering of. Hierarchical clustering divides the reactions into multiple clusters depending on a tunable level of clustering. On one end, all reactions are clubbed into a single cluster, and on the other end, every reaction is its own cluster. For any given level of clustering, we employed a simple rule to predict missing reactions from known reactions. If any one of the reactions in a cluster is known to be present in the metabolic repertoire of a bacterium, all other reactions in the cluster are also predicted to be in the network. Using this simple prediction model and by varying the level of clustering, we obtained true and false positive rates for the predictions for each bacteria. These rates were averaged across all bacteria for a given level of clustering and plotted as a receiver operating characteristic (ROC) curve. Figure 4a shows that this simple approach to predict the missing metabolic reactions performs exceedingly well. The area under the curve when only 10% of the reactions are known is 0.916, which increases to 0.988 when 90% of the reactions are known. Notably, the performance of our approach is extremely accurate across different values of *K* and different fractions of missing reactions (SI Figure 3). Importantly, the highest overlap between the true metabolic repertoire and the predicted metabolic repertoire as quantified by the Jaccard index occurs near the true size of the network (Fig. 4b).

**Figure 4.**
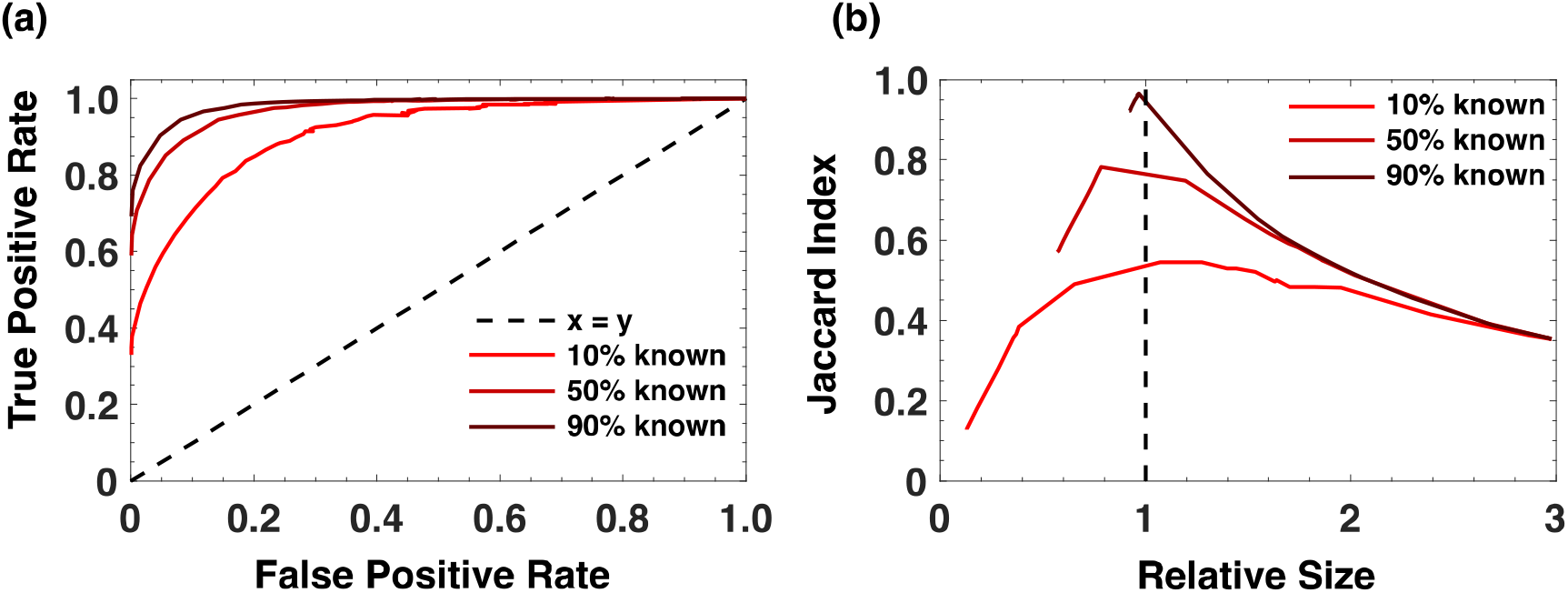
SiGMoiD predicts the presence/absence of metabolic reactions. **(a)** Receiver operating characteristic (ROC) curve for SiGMoiD-based prediction of missing metabolic reactions. Different lines represent metabolic models with different fractions of known reactions. **(b)** The mean Jaccard index between the set of predicted metabolic reactions and the actual metabolic reactions in any species (y-axis) vs. the relative size of the predicted network to the actual network (x-axis).

Notably, unlike other gene inference methods that rely on phylogenetic placement of the query genome against a reference database^26,27^ or approaches that can use phenotypic knowledge about the organism (nutrients that can sustain its growth, biomass composition, etc.)^28^ our approach predicts reactions solely based on their co-occurrence profile across the bacterial kingdom. Therefore, our approach can be integrated with other methods to robustly predict missing reactions in genome scale metabolic reconstructions.

## Discussion

A deluge of biophysical data in the last decade has called for the development of top-down modeling approaches. Here, instead of describing the data from first principles mechanistic models, one constructs probability distributions that represent it. As a result, generative models of collective behavior have become essential to modeling several biophysical systems. The most popular way to generate top-down models is the maximum entropy (max ent) approach wherein one approximates the data using a probabilistic model that reproduces lower order statistics estimated from the data. The max ent approach has the significant conceptual advantage that it represents the simplest model consistent with the imposed constraints. However, there are two significant drawbacks. First, the constraints are hand-picked by the modeler and the model therefore depends on these constraints. For binary data, constraints of averages and pair correlations have become popular. Second, the inference of max ent models for large data sets can be computationally expensive and it may be unrealistic to infer models for > 100 binary variables.

To address these issues, we developed SiGMoiD. SiGMoiD takes an agnostic approach about the constraints. In SiGMoiD, instead of specifying the constraints, the user only specifies the total number of constraints. SiGMoiD learns these constraints from the data. Moreover, parameter inference in SiGMoiD is orders of magnitude faster than max ent inference. We showed using three data sets of varying complexity that SiGMoiD not only performs as well as max ent models in terms of accuracy but can also be applied to study very large data sets that are currently out of the reach of max ent inference. Going forward, we believe that this computationally efficient and conceptually straightforward approach will be immensely valuable in modeling collective behavior of high dimensional data.

## Acknowledgments

XZ and PD would like to thank the UF startup fund.

## Supplementary Information

### Curating the data

All the data used in this study and the code used to analyze the data can be found on github: https://github.com/zhaoxc099/sigmoid

### Collective firing of neurons

Neuron firing data was obtained from an open source repository^13,14^. We combined the data across all trials and all time points and selected 15 neurons that had the highest overall firing propensity. We randomly selected 10^4^ samples from the data for further analysis.

### Co-occurrence of bacterial species

Bacterial co-occurrence data was downloaded as an OTU table from Sheth et al.^3^. The OTU table was binarized by assigning a 1 if a particular OTU was present (positive abundance) in any given sample and zero otherwise. We removed the OTUs that were present in none of the samples from our analysis.

### Bacterial metabolic networks

From a list of ∼ 27000 annotated bacterial genomes in the base database^24^ (March 2017) a representative genome was selected for each named species. The representative genome was chosen so that it was close to the median number of genes across all genomes for the corresponding species. Genomes that were obtained using single-cell sequencing were not considered due to the possibility of low genome coverage. For each selected genome, we built a draft genome-scale metabolic reconstruction using the ‘Build Metabolic Model’ method (v.1.5.1) in the kbase Narrative interface. The method implements the ModelSeed pipeline described by Henry et al.^25^. We only considered reconstructions with less than 30% reactions added during the gap-filling step of the ModelSeed pipeline as a proxy for high quality models.

The gap-filling approach taken by Kbase involves minimization of an arbitrarily trained optimization function. Therefore, in our analysis we only included reactions that were annotated to a gene or to a combination of genes. At the end, we had a total of ∼ 4000 bacterial metabolic models with a metabolic universe of ∼ 3300 reactions. 400 models were randomly chosen to be in the testing data set and the rest were used to infer SiGMoiD model parameters.

## Supplementary Figures

**SI Figure 1.**
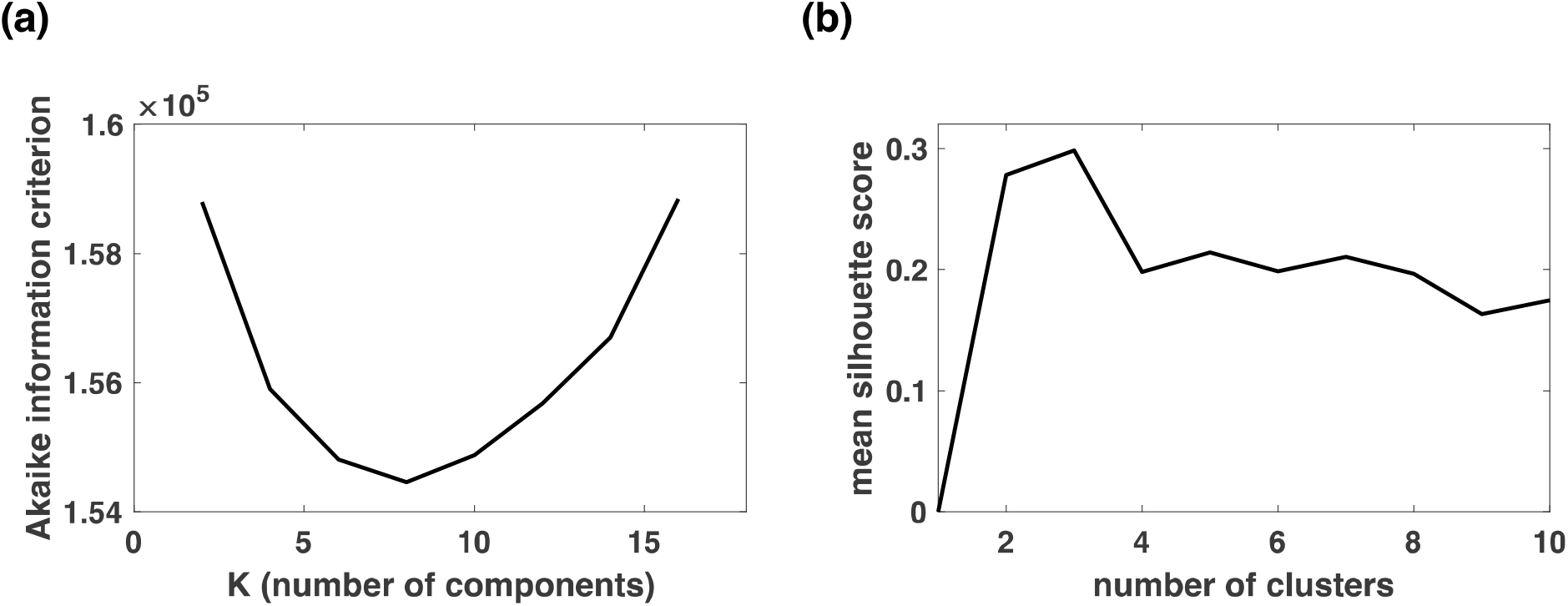
**(a)** Akaike information criterion as a function of K, the number of components used to model the microbiome co-occurrence data. **(b)** Mean silhouette score as a number of clusters using K-means clustering of the particle-specific ***β***_s_ vectors.

**SI Figure 2.**
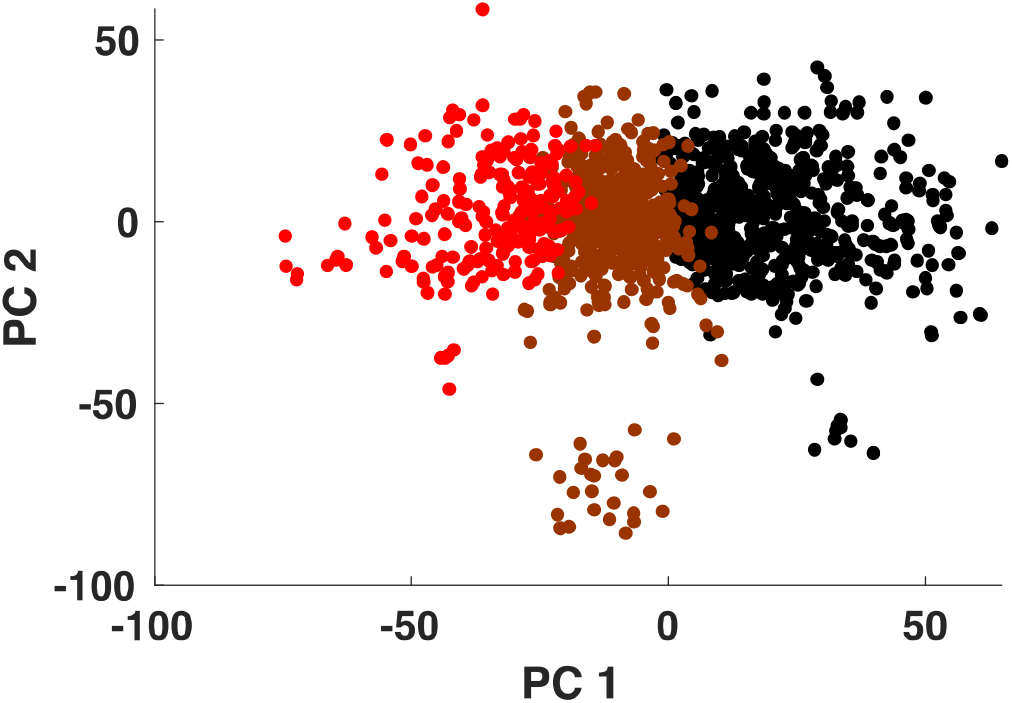
n=3 clusters of particles were identified using K-means clustering of the microbiome co-occurrence, shown here using the first two principal components of the particle-specific ***β***_s_ vectors.

**SI Figure 3.**
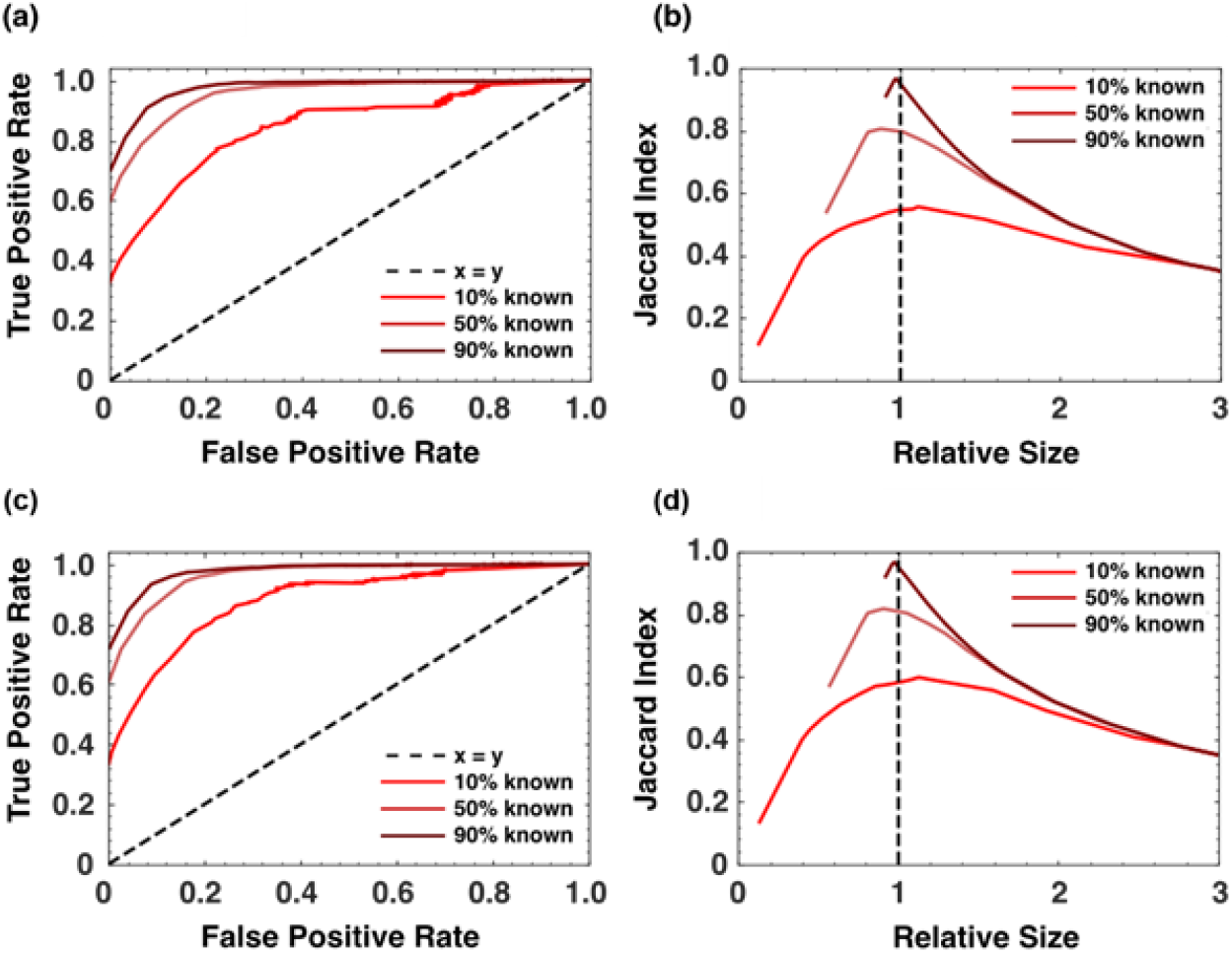
Performance of the SiGMoiD-based approach to identify missing metabolic reactions with K = 20 components (panels a and b) and K = 40 components (panels c and d).

## Supplementary Tables

**SI Table 1.**
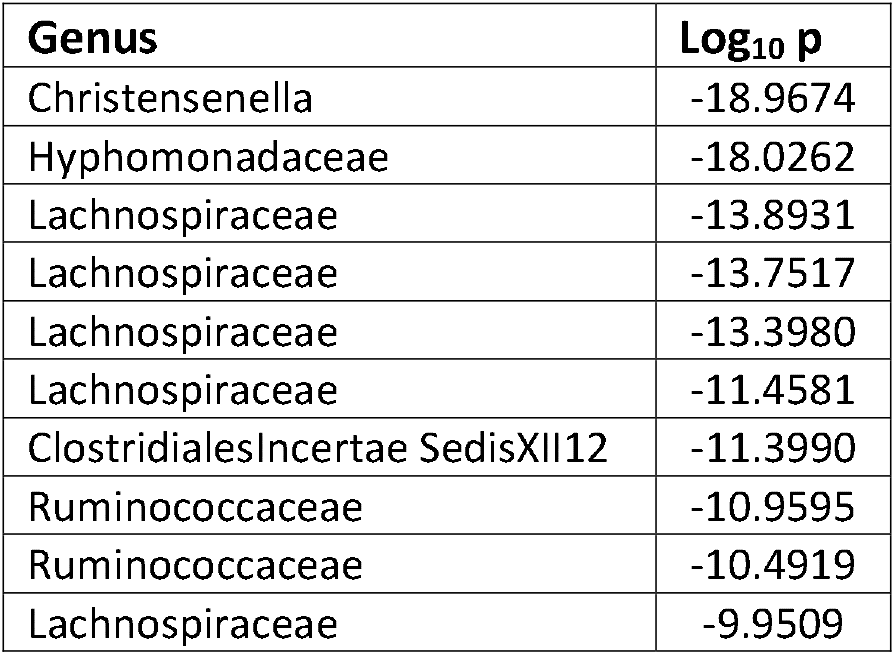
Genera of OTUs that are most enriched in cluster 3 using a hypergeometric test and the corresponding p-values.

